# Genotypic differences in wheat yield determinants within a NAM population based on elite parents

**DOI:** 10.1101/2020.09.08.287763

**Authors:** Priyanka A. Basavaraddi, Roxana Savin, Sivakumar Sukumaran, Matthew P. Reynolds, Simon Griffiths, Gustavo A. Slafer

**Affiliations:** Department of Crop and Forest Sciences, University of Lleida - AGROTECNIO Center, Av. R. Roure 191, 25198 Lleida, Spain; International Maize and Wheat Improvement Center (CIMMYT), El Batán, Texcoco CP 56237, Mexico; John Innes Centre, Norwich Research Park, Colney Ln, Norwich, NR4 7UH, United Kingdom; ICREA, Catalonian Institution for Research and Advanced Studies, Spain

**Keywords:** Pre-anthesis phases, fruiting efficiency, spike dry weight at anthesis, source-sink, yield components

## Abstract

Future grain yield (GY) improvements require the identification of beneficial traits within the context of high yield potential and not just based on the pleiotropic effect of traits such as crop height and heading date. We evaluated 1937 lines from Nested Association Mapping (NAM) population derived from 13 bi-parental varietal crosses under field conditions. We selected 493 lines with similar time to anthesis to that of the two checks used in the study (across and within each family) which reduced the range of plant height in the selected lines. Yield components were measured in these 493 lines from which 231 lines were selected by excluding lines with lowest number of grains so excluded low yielding lines. Later the subset of 231 lines were evaluated in two field experiments (2016-17, CS1 and 2017-18, CS2). Numerical and physiological components of grain yield were measured. The two-step selection maximised GY within an acceptable range of variation for height and anthesis. GY in 231 lines showed very high G×E interaction. Taking both seasons together, we selected lines from upper and lower quartile GY groups to identify stable beneficial trait combinations for improved GY. Differences in GY were explained by grain number driven by increased spike dry weight at anthesis (SDWa) and fruiting efficiency (FE). Increased GY was accompanied by sink limitation. The data points towards increases in grain number as the route towards future GY increases in wheat breeding.

## 1. Introduction

Present rates of genetic gains in wheat grain yield (GY) are insufficient to satisfy future demands (Reynolds et al., 2012) which is estimated to increase 50% by 2050 from current level of demand (https://www.cimmyt.org/work/wheat-research/; accessed on 05.05.2020). In recent decades the rate of genetic gain has decreased (e.g. Aisawi et al., 2015; Flohr et al., 2018; Maeoka et al., 2020), in many cases to a standstill (e.g. Acreche et al., 2008; Chairi et al., 2018; de Oliveira Silva et al., 2020; Lo Valvo et al., 2018). To address this problem, we need to improve our understanding of physiological attributes likely to underpin future GY gains as well as to identify variation available within elite germplasm for these traits. Grain number per m^2^ (GN) and average grain weight (AGW) are the two most important GY components (Slafer et al., 2014). Owing to larger plasticity it is GN that has delivered most GY improvements (Abbate et al., 1995; Calderini and Slafer, 1999; Fischer, 1985; Reynolds et al., 2009; Serrago et al., 2013; Siddique et al., 1989a; Slafer et al., 1990; Slafer and Andrade, 1989), even though it has much lower heritability than AGW (Sadras and Slafer, 2012).

Past improvements in wheat GN and GY came through the gradual accumulation of beneficial quantitative variation as well as a limited set of step changes such as the widespread deployment of semi-dwarf genes, chiefly Rht-1 (e.g. Calderini and Slafer, 1999; Flintham et al., 1997) and improving adaptation by changing time to anthesis to be more adequate for a specific region (e.g. Araus et al., 2002) particularly through changes in photoperiod and vernalisation sensitivity (González et al., 2005a; Griffiths et al., 2009; Shaw et al., 2012; Whitechurch and Snape, 2003). Reductions in plant height mediated by *R*ht-1 enhanced biomass partitioning to the juvenile spikes prior to anthesis (Brooking and Kirby, 1981; Fischer and Stockman, 1986; Miralles et al., 1998) which in turn allowed for an improved development of florets resulting in higher GN (Ferrante et al., 2013; Fischer and Stockman, 1986; Miralles et al., 1998; Siddique et al., 1989a). Introgression of Rht-1 alleles and homoeoalleles increased harvest index (HI) through increased GN and improved GY without major changes in biomass and a reduction in AGW, that naturally did not counteract the GN benefits (Bingham and Wellington., 1981; Calderini et al., 1995; Flintham et al., 1997; Miralles and Slafer, 1995; Shearman et al., 2005; Siddique et al., 1989b). Adjustments in time to anthesis have been critical to improve GY through improving adaptation mainly when the life cycle of the original genotypes exploited in a region did not allow maximum use of available resources or for stress avoidance (Araus et al., 2002). These two traits, that have been critical to improve yields in the past, would be of limited importance in the future as they have already been optimised in major wheat growing regions (e.g. Acreche et al., 2008; Maeoka et al., 2020; Slafer et al., 2005).

Future gains in GN will provide the increased sink strength which many studies have pointed to as required to increase GY, because of the frequent sink limitation for grain filling in wheat (Borrás et al., 2004; Borrill et al., 2015; Reynolds et al., 2005; Serrago et al., 2013 and referennces quoted there in). Final GN is a highly integrative trait (highly plastic and with low heritability; Sadras and Slafer, 2012b) with many of the development processes that lead to contributing to the final number. So, the identification of major genes or QTL directly and consistently controlling it is unlikely. For these reasons it is important to understand which traits are responsible for differences in GN within elite material and to show how they could be deployed by breeders aiming to improve GY within elite × elite pedigrees by reducing sink limitation in their finished varieties.

While time to anthesis is tightly controlled in breeders selections around a local optimum, the partitioning of the cycle into different duration of phases occurring before and after terminal spikelet (TS) might still be improved (Slafer et al., 2001). Components of GN are formed from sowing to a few days after anthesis (Slafer and Rawson, 1994) but the most sensitive phase is demarcated by TS and anthesis, the late reproductive phase or LRP (Slafer, 2003; Fischer, 2011), and in particular the last half of it. Thus, it has been hypothesised that lengthening the duration of the LRP, when floret development takes place, would improve GN (Miralles and Slafer, 2007).

By the time anthesis is reached the stage is set for the realisation of GN, in fact the spike dry weight at anthesis (SDWa) has been shown to be highly predictive of GN in a number of experiments (Fischer, 2011; Ferrante et al., 2013). The physiological support for this mechanistic relationship is that floret primordia survival is closely linked to SDWa and in wheat, being a cleistogamous plant, most fertile florets become grains after anthesis. The number of fertile florets at anthesis depends mainly on the balance between the initiation and mortality of floret primordia during the LRP (Kirby, 1988; Prieto et al., 2018). Both floret mortality (Ferrante et al., 2013; González et al., 2011) and survival (Ferrante et al., 2013, 2012; González et al., 2005b; Siddique et al., 1989a) seems to depend on the availability of resources for spike growth from flag leaf appearance to anthesis. The physiological dissection of this point in development has been taken further by Slafer *et al*. (2015) using the concept of fruiting efficiency (FE, number of grains produced per unit SDWa) and showing that FE can be useful towards genetically improving wheat GY (see also empirical proofs in Acreche et al., 2008; Flohr et al., 2018; Lo Valvo et al., 2018b; Zhang et al., 2019).

For a proper identification of traits or trait combinations that are likely to be important and useful in modern wheat breeding programmes (i.e. beyond traits like plant height which are already optimised), it is important to study trait relationships within the context of elite germplasm. Although the analyses restricted to elite genotypes will naturally reduce substantially the degree of variation that could be expected from unselected lines of wider crosses (and would consequently yield less clear relationships). The advantage is that the materials used would resemble better what realistic breeding does (crosses of elite × elite) when aiming to improve yield, and therefore results and conclusions would be more likely truly applicable in actual breeding programmes. Therefore, in the present study we firstly grew a very large population of elite lines (1937 lines of a Nested Association Mapping, NAM, population produced by crossing elite parents) in the field at Ciudad Obregón, Mexico (Cd. Obregón) and from these initial results we further selected a relatively small, yet rather large, sub-set of 231 lines that were considered best performing (within germplasm that was already elite) to study them more in detail in field experiments carried out in Bell-lloc d’Urgell, Spain (Bell-lloc) over two cropping seasons.

## 2. Materials and Methods

### 2.1. Experimental field conditions

The first field evaluation of the whole NAM population was carried out in the 2015-16 cropping season at CIMMYT’s experimental station (within the Norman E. Borlaug Experimental Field, CENEB) in Cd. Obregón, Sonora, North-West Mexico (lat. 27°23’ N, 109°55’W). The experiment was sown on 10 December 2015 in small plots (“hills”, 80 cm between hills, 30 cm long) at a density equivalent to 5 plants per plot.

In the following two seasons (2016-17, CS1 and 2017-18, CS2), field experiments were carried out near Bell-lloc d’Urgell, Lleida, North-East Spain (Lat. 41°38′ N, 0°44′ E in CS1 and Lat. 41°37′ N, 0°47′ E in CS2). Experiments were sown on 16 November 2016 and on 17 November 2017, both at the rate of 125 kg ha^−1^ aiming to attain an effective plant density of 250 plants per m^2^. The three experiments were carried out avoiding stresses: plots were always sown within the optimal dates to maximize yield, fully fertilized, irrigated, Weeds, pests and diseases were prevented or controlled. Soil nitrogen availability was determined in CS1 and CS2 at the beginning of the experiments. Eight samples from the soil surface to 0.9 m depth were randomly taken from the field were the experiments were sown and analysed for mineral N content. The average available N content of the experimental area was 133.1±9.3 and 115.4±8.8 KgN ha^−1^ in CS1 and CS2, respectively. This soil nitrogen availability was supplemented with 150 KgN ha^−1^ (as urea) uniformly applied to each plot at the onset of tillering.

Meteorological data for the cropping periods were recorded from the Meteorological station located near CENEB for the first experiment and from the Meteorological station of Meteocat (Servei Meteorologic de Catalunya) close to the experimental fields in the last two experiments (Table 1).

**Table 1:**
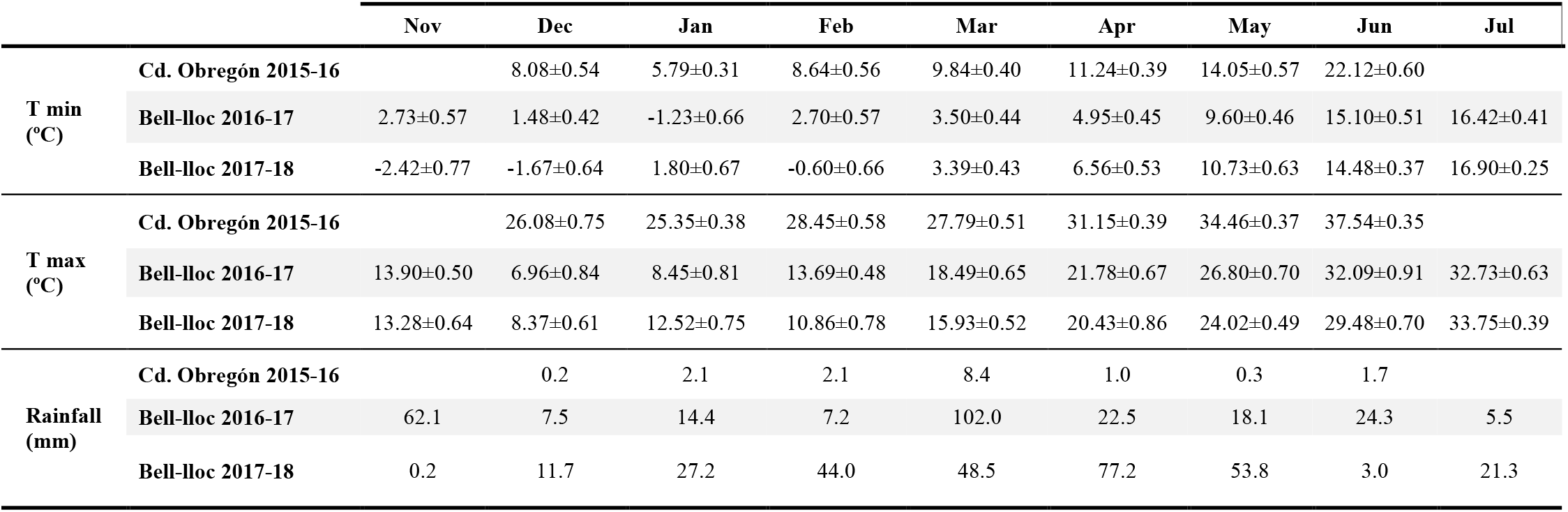
Meteorological data for experiments in Ciudad Obregón 2015-16, and in Bell-lloc 2016-17 (CS 1) and 2017-18 (CS 2): monthly average of minimum (T min) and maximum (T max) temperatures (± standard error) as well as monthly cumulative precipitation. In all cases data are provided for the growing season (from the month of sowing to that of harvest).

In Bell-lloc (where the more detailed experiments were carried out), the average temperature for the whole cropping duration (November to July) of CS1 was 12.6 °C whereas CS2 was 11.7 °C. At the critical stage of anthesis both minimum and maximum temperatures were slightly higher during CS1 than CS2. In general, temperatures, both minimum and maximum were within the ranges normally occurring in the region during past 5 years. As mentioned above, the experimental fields were irrigated as needed: in Bell-lloc both experiments were irrigated around anthesis but in CS2 an additional irrigation was given at seedling emergence stage as late fall – early winter of 2017 was unusually dry (Table 1). Thus, there was only one irrigation in CS1 (on 19 April 2017) and two in CS2 (on 14 December 2017 and 5 May 2018). Each irrigation was equivalent to 80 mm of rainfall.

### 2.2. Genotypes and experimental design

In the experiment at Cd. Obregón we grew the whole NAM population while in the two field experiments at Bell-lloc we grew a selection of this population. The NAM population was generated from 13 bi-parental crosses where both parents in each cross were elite spring wheat varieties selected for having particular traits of interest to be included in the crosses. The parents of the crosses used were (i) Paragon, one of the best UK spring wheat cultivars considering yield potential and disease resistance, was the most common parent used; (ii) four CIMCOG (CIMMYT Core Germplasm: Orford et al., 2014) lines viz.: CIMCOG 49, 47, 3 and 32 characterised for their high values of biomass, grains per spike and harvest index; (iii) Weebill, a cultivar well known for having its high yield associated to superior average grain weight; (iv) MISR1, SUPER152, Pfau, Waxwing and Baj, all parents selected for their high yield related to earliness in time to anthesis; and (v) Wyalkatchem a high performing Australian variety. The 13 families were the lines derived of the following crosses: (1) Weebill × CIMCOG3, (2) Weebill × CIMCOG32, (3) Paragon × Pfau, (4) Paragon × Baj, (5) Paragon × Wyalkatchem, (6) Paragon × (Becard × Kachu), (7) Paragon × MISR1, (8) Paragon × Waxwing, (9) Paragon × Garcia, (10) Paragon × Super151, (11) Paragon × Synth type, (12) Paragon × CIMCOG47, and (13) Paragon × CIMCOG49 (please note that the order of the crosses mentioned here from 1 to 13 will be followed in the result section). Detailed descriptions of the populations, including Axiom 35K genotype files and genetic maps can be found at https://data.cimmyt.org/dataset.xhtml?persistentId=hdl:11529/10996, accessed on 05.05.2020. All germplasm is deposited at the CIMMYT genebank.

In the experiment at Cd. Obregón, the original set of 1937 lines were grown in un-replicated hill-plots together with checks (the parents of the crosses and two well adapted genotypes, Reedling and Sokoll) replicated across the whole experiment. Plots were arranged as different families with embedded checks in an augmented design (considering the lines of the NAM, parents and replicated checks there were 2120 hill plots).

In the two experiments conducted in Bell-lloc, we grew 231 lines which is a sub-set from 1937 lines selected based on their field performances in the initial experiment at Cd. Obregón. Treatments consisted of 231 selected lines grown in un-replicated plots together with replicated check plots across the experiments using augmented randomised complete block design in a regular grid, design which is commonly used to test large populations where it is not possible to have a complete replication of lines (Scott and Milliken, 1993). Plots in both experiments were arranged in the field with random allocation of un-replicated 231 genotypes and replicated 3 checks (there were 26 check plots arranged in order to have two check plots in each of the 13 rows of plots arranged diagonally across rows of plots; Müller et al., 2010). The layout of the experiments had 13 rows and 20 columns of plots making it a total of 260 plots, of which 257 corresponded to lines and checks in which traits were measured (the other three plots were sown to complete the rectangular field layout but were not measured). In addition, the whole experiment had a set of 70 border plots that were not considered for measurements (were sown and maintained to avoid border effects on the plots allocated to rows 1 and 13 and to columns 1 and 20 of the measured plots). Each plot consisting of 6 rows was 0.2 m apart and 4 m long. Three cultivars viz., Paragon, Garcia and Paledor were the checks used both to quantify the spatial heterogeneity across field and as a reference for performance of well adapted cultivars. Paragon was used, as it was the most common parent of the studied NAM population while Garcia (http://www.genvce.org/variedades/trigo-blando/invierno/garcia/; accessed on 14.01.2020 or http://www.agrusa.com/Semillas.php?_b=&_un=1&_do=18&_tr=19; accessed on 05.05.2020) and Paledor (http://www.genvce.org/variedades/trigo-blando/invierno/paledor/; accessed on 14.01.2020 and http://www.agrusa.com/Semillas.php?_b=&_un=1&_do=18&_tr=22; accessed on 05.05.2020) were chosen to be two of the best performing local cultivars at the time we conducted the study. Paledor was indeed a check in the variety trials at least until the cropping season immediately before the CS1 (https://genvce.org/wp-content/uploads/2019/12/informe-genvce-cereal-de-invierno-2015-2016.pdf, accessed on 27.08.2020).

### 2.3. Measurements and determinations

In the field experiment conducted at Cd. Obregón plant height and anthesis date were determined in all the 2120 hill plots. Based on these determinations, 493 lines, which had similar time to anthesis to that of the checks and discarding extremely short lines, were sampled at maturity and yield per hill as well as AGW were determined. Of these 493 lines, the 231 lines that exhibited best field performance were selected to be evaluated in the more detailed study carried out over the following two seasons in Bell-lloc.

In the two field experiments in Bell-lloc we determined in each plot different stages of development using the decimal code developed by Zadoks et al. (1974): seedling emergence (stage DC10), onset of stem elongation (DC30), flag leaf emergence (DC39), heading (DC59), anthesis (DC65) and physiological maturity (DC95). All the stages were recorded when 50% of the plot showed that stage by monitoring each plot regularly (from once a week to thrice a week, depending on temperature). The onset of stem elongation (OSE) was determined by touching the main shoot at the base just above the ground to detect the first node and was repeated on several plants in each plot to record the stage for that plot. Later, the OSE data from a parallel but smaller experiment conducted in the same field, in which we also determined the stage of TS by periodic dissection of the apex, was used to estimate the timing of TS from the OSE measurements. Length of phenological phases was estimated in thermal time with base temperature of 0 °C.

Plants were sampled at anthesis (stage DC65) and physiological maturity (DC95) from each individual plot from 1 linear meter which was chosen randomly (from any of the 4 central rows and avoiding the extreme 25 cm of the rows that were left as borders). Plants in that sampling area were manually pulled out to recover the whole above ground biomass and taken to the laboratory where they were processed to record number of plants, shoots, and productive shoots (shoots bearing spikes) and stem length from the soil level to the base of the spikes. Leaves (only leaf laminae), spikes and stems (including leaf sheaths) were separated and dried in a hot-air oven at 65 °C for 72 h after which dry weights were recorded. At physiological maturity, spikes were dried and weighed and were threshed to obtain grains. Later, the grains were counted and dried again for at least 24 h to measure the grain weights.

### 2.4. Analyses

For the data from field experiment in Cd. Obregón only descriptive statistics were performed. Data from the field experiments in Bell-lloc were analysed using GENSTAT Pro (Version 19) in the preliminary single environment analysis where checks are considered as extra genotypes that are replicated and effect of spatial heterogeneity on un-replicated lines were accounted for using variation observed in checks which is then used to estimate Best Linear Unbiased Estimates (BLUEs). Relationships between traits were analysed with linear regressions.

## 3. Results

### 3.1. Genetic variation in the whole NAM population and selection of a sub-set

Expectedly, the ranges of variation in phenology and in plant height were rather large when considering the whole NAM population of 1937 lines (Fig. 1).

**Figure 1.**
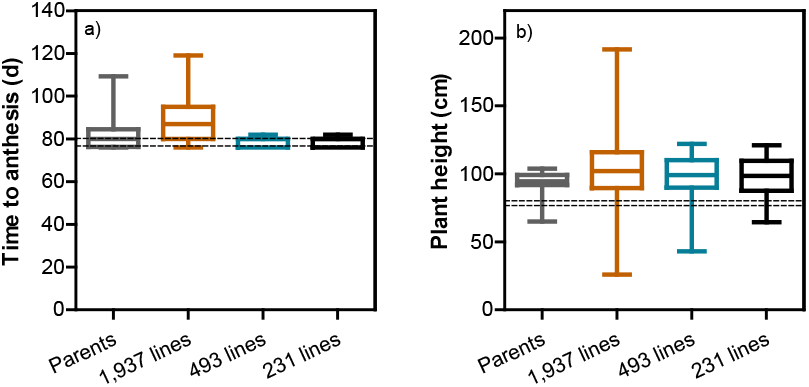
Boxplots for time from sowing to anthesis (a) and plant height (b) from the experiment carried out in Cd. Obregón (NW Mexico) in 2015-16 considering the variability within the parents of the 13 crosses, the whole original NAM population of 1,937 lines, the 493 lines that were sampled and for which yield components were determined, and the 231 lines that were finally selected to be further studied in later field experiments carried out in Bell-lloc (NE Spain). Dashed lines show the values corresponding to two well adapted genotypes used as checks in the experiment, viz. Reedling and Sokoll.

Time to anthesis ranged from c. 75 to c. 120 d (Fig. 1a), equivalent to a thermal time range from c. 1150 to c. 2000°C d with a base temperature of 0° C. This large degree of variation was due to a few parents that had a considerably longer time to anthesis than most others in Cd Obregón as well as a large transgressive segregation particularly for longer periods to anthesis, as the longest times to anthesis in the lines analysed exceeded, by c. 10 d (c. 212°C d), the already large range of variation shown by the parents of the population (Fig. 1a). This was in part due to the inclusion of cultivars possessing valuable yield-determining traits beyond time to anthesis (such as Paragon) with a strong photoperiod sensitivity conferring late flowering and maladaptation in Cd. Obregón though many of the lines derived from Paragon would (c. three quarter of the parents differed in time to anthesis by less than a week in this growing condition and half of them flowered within the two days of difference shown by the two well adapted genotypes used as checks; Fig. 1a). As we aimed to identify traits of value beyond time to anthesis and plant height, the data from the first experiment was used to select against variation in time to anthesis that exceeded that of the best adapted local check varieties. Thus, the range of variation in time to anthesis in the selected 493 lines (which were then sampled at physiological maturity to measure hill-plot yield and AGW) was dramatically reduced (Fig. 1a), and could not be further reduced when selecting the 231 lines for later experiments (Fig. 1a).

Plant height in the whole NAM population also varied hugely, from c. 0.25 to almost 2 m (Fig. 1b) and in this case mostly due to large transgressive segregation (likely due to segregation of Rht alleles resulting in some lines being tall and others double dwarf), as parents of the 13 crosses ranged in height from c. 0.6 to 1.1 m and most parents had a height very similar to that of the two well adapted checks (Fig. 1b). The selection of lines that had time to anthesis in the narrow range of best adapted checks reduced the range of variability in height to c. 0.5 to 1.2 m and the final selection of lines to be further tested in later experiments reduced that variation further by discarding the shortest plants (<0.64 m; Fig. 1b).

To produce the final selection of the subset of 231 lines to be analysed in more detail we considered the hill-plot yield and yield components of the 493 lines that were sampled in the experiment. The range of GY and its two major components were relatively large (Fig. 2), even though these lines displayed virtually no difference in time to anthesis and exhibited a range of plant height that is substantially reduced compared to the whole NAM population. Indeed, variation in time to anthesis or in plant height explained a negligible proportion of the genotypic variation in GY among the 493 lines (0.4% and 6.6%, respectively). As GY was more related to the number than to the weight of grains, and these major components were negatively related to each other (Supplementary Fig. S1), the selection was made eliminating the lines with lowest number of grains and lowest GY. Consequently, the selected sub-set of 231 lines (17-18 lines per each of the 13 crosses) reduced the variability in GY and its two components shown in the set of 493 lines, through maintaining the lines with highest GY and GN per hill as well as with intermediate values of AGW (Fig. 2).

**Figure 2.**
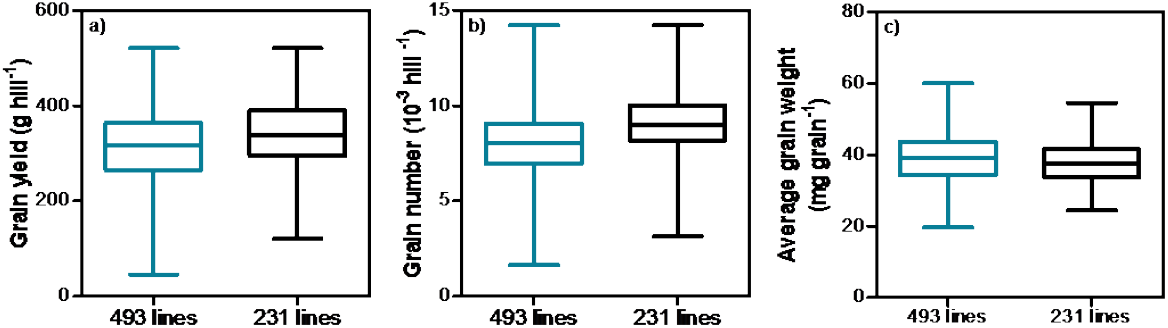
Boxplots of grain yield (a) and its two major components, grain number (b) and their average weight (c) in the experiment carried out in Cd. Obregón (NW Mexico) in 2015-16 considering the variability for the 493 lines that had similar time to anthesis to that of the well adapted checks, and the 231 lines that were further selected to vary less in plant height and which were finally selected to be further studied in later field experiments carried out in Bell-lloc (NE Spain).

### 3.2. Genetic variation in, and relationships between, GY and phenology within the selected lines

When in the next two seasons these selected 231 lines were grown in Bell-lloc, the range of variation in time to anthesis was larger than for the same lines which had been selected in Cd. Obregón with the aim of constraining anthesis data. However, the timeframe of anthesis was much narrower than would be expected from the whole NAM (n=1937) or a random selection of it. Even though the length of cropping cycle is longer in Spain than Mexico, the range in time to anthesis for the selected panel of 231 lines was much narrower than the original population of 1937 lines evaluated in Cd. Obregón (cf. Figs. 1a and 3a and b). It is also true that although lines were selected for having similar time to anthesis within the families, there was a noticeable variation, not only considering the whole population (1188-1481 °C d in CS1 and 1209-1525 °C d in CS2) but also within most families (Fig. 3a, b). There was a reasonable degree of consistency for thermal time to anthesis between the two cropping seasons (Fig. 3c).

**Figure 3.**
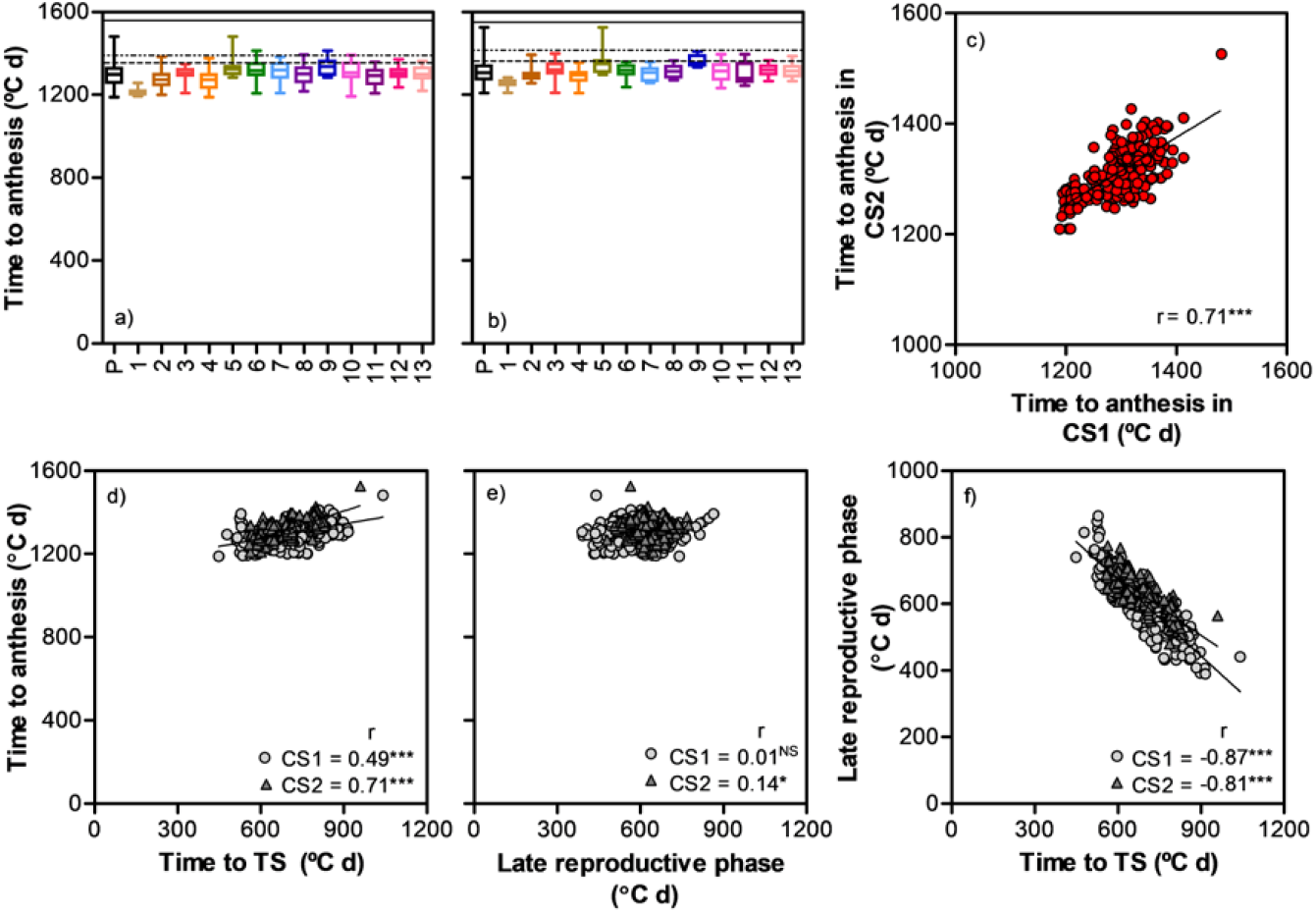
Upper panels: Boxplots showing variability in time from sowing to anthesis within the whole population (P) and within families (13 bi-parental crosses) along with three checks Paragon (solid line), Paledor (dotted line) and Garcia (dashed line) in the first (CS1, a) and second cropping season (CS2, b), and consistency of time to anthesis over the two cropping seasons (c). Bottom panels: Relationships between time to anthesis and its component phases: time from sowing to terminal spikelet (TS, d) and time from then to anthesis, i.e. the late reproductive phase (e); as well as between the two component phases (f) for the 231 lines grown in the first (CS1) and second (CS2) cropping seasons. Note: The crosses corresponding to serial number 1-13 is given in materials and methods; graphs c, d, e and f do not include checks; origin of the graph c does not begin at 0. Coefficients of correlations and their significance level (* p<0.05; *** p<0.001; NS=non-significant) are shown for the relationships.

The same was true for plant height: lines differed in height both across and within families but genotypic differences in height were reasonably consistent across both seasons (Supplementary Fig.S2).

Genetic variation in thermal time to anthesis was related to variation for each of the two component phases considered: time from sowing to TS (Fig. 3d) and time since then to anthesis (Fig. 3e), though the correlations were stronger with the initial phase to TS, embracing the vegetative and early reproductive phases, than with the LRP, suggesting that variation in time to anthesis was mainly controlled by the duration of the first phase in this panel. Furthermore, there was a clear trend for compensation between duration of these two phases both the seasons (Fig. 3f), that was naturally only partial (otherwise there would have been no variation in thermal time to anthesis), as revealed by the slopes that were higher (i.e. less negative) than −1 (−0.76 and −0.57 in CS1 and CS2, respectively). This means that in both cropping seasons it was possible to identify lines with the same time to anthesis but differing oppositely in the duration of the phases of leaf and spikelet initiation and of floret development.

Genotypic variation in GY was large (440 to 1181 g m^−2^ and 459 to 1067 g m^−2^ in CS1 and CS2, respectively). And once again, the variation across the subset of the NAM population reflected more the variation within families than differences across families (Fig. 4a, b). For most of the families, and therefore for the whole population, there were several lines with greater GY than the local checks, Paledor and Garcia, which were modern commercial high-yielding cultivars. However, unlike with time to anthesis and plant height, there was a very large G×E interaction for GY, evident from the absence of significant relationship between GY of the lines across the two cropping seasons (Fig. 4c). This lack of consistency between seasons was also evident for the yield difference between the two well adapted cultivars: while their difference was not significant in CS1 (7.82±0.40 and 7.12±0.42 Mg ha-1) it was larger and highly significant in CS2 (9.19±0.46 and 7.70±0.17).

**Figure 4.**
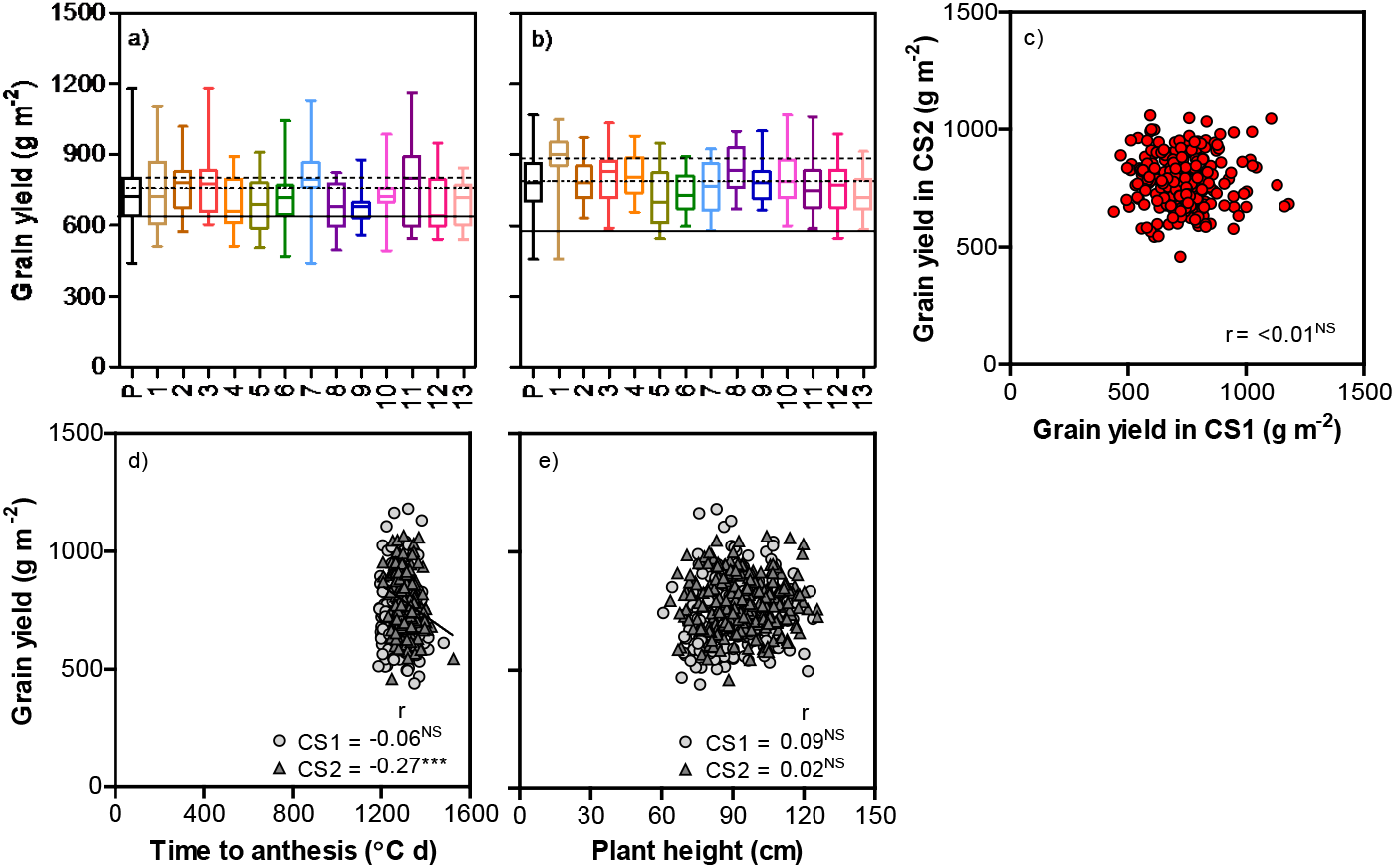
Upper panels: Boxplots showing large variability for grain yield within the whole population (P) and within each family (13 bi-parental crosses) along with three checks Paragon (solid line), Paledor (dotted line) and Garcia (dashed line) in the first (CS1, a) and second cropping season (CS2, b) and inconsistency for grain yield over two seasons (c). Bottom panels: Relationships between grain yield and either total time to anthesis (d) or plant height (e). Note: The crosses corresponding to serial number 1-13 is given in materials and methods; graph c, d and e do not include checks. Coefficients of correlations and their significance level (*** p<0.001; NS=non-significant) are shown for the relationships.

Although the differences in time to anthesis and plant height could potentially interfere with the relationships between GY and traits other than these two, such potential interference would not be critical in this study as there were no clear relationships between GY and either time to anthesis (Fig. 4d) or plant height (Fig. 4e). Although time to anthesis was significantly related to GY in CS2, it explained only 7% of the GY variation, whilst time to anthesis in CS1 and plant height in both seasons explained less than 1% of the variation in GY (Fig. 4d, e).

Even though the total time to anthesis did not explain differences in GY within the whole population, the partitioning of that time into phases occurring before or after TS seemed to have some significance: across lines and within each of the two seasons GY tended to be negatively related to the duration of the first phase, time from sowing to TS (Supplementary Fig. S3a) and positively related to the length of the following phase, the LRP (Supplementary Fig. S3b). However, even when statistically significant, these relations were weak.

Taking into account the large G×E interaction (reflected by the lack of consistency in GY between CS1 and CS2) and our aim to identify trait relationships that can suggest traits relevant for further raising yield through breeding, we identified sub-groups within the sub-set of 231 lines that consistently expressed the extremes of GY over the two seasons: we chose all lines that were part of the bottom and top quartiles of GY in both seasons, low- and high-GY (GY_L_ and GY_H_, respectively). Applying this criterion, there were 13 GY_L_ and 11 GY_H_ lines.

Naturally these two sub-groups of lines differed in GY highly significantly, with no overlap between them (i.e. the lowest-yielding line of GY_H_ clearly out yielded the highest-yielding line of GY_L_; Fig. 5). However, there were also clear genotypic differences in GY within each of these two sub-groups (Fig. 5). By virtue of the data selection made, major genetic variation in GY between the two sub-groups were highly consistent across seasons. We focused on the analysis of traits determining GY in the selected lines within and across these GY_L_ and GY_H_. For the benefit of readers who may be interested in the relationships across the whole sub-set of 231 lines we did also report the relationships for them, naturally for each season separately due to the large G×E interaction in GY, in supplementary figures. The mainstream relationships will be shown both for the two sub-groups separately (highlighting whether the considered trait was relevant or not for the genetic variation within GY_L_ and GY_H_ lines) and for all of them together (highlighting the contribution of the trait to produce the consistently highest-yielding lines of the population) and described the variation levels in these traits between these two sub-groups.

**Figure 5.**
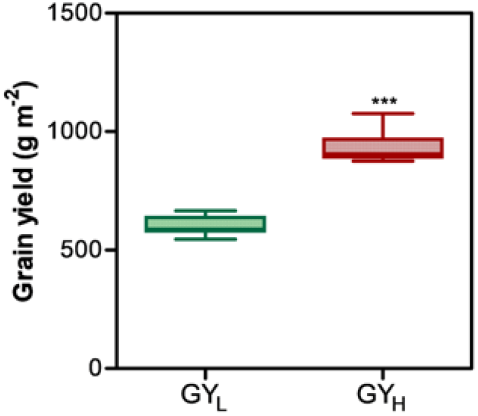
Boxplot depicting variation in grain yield between selected sub-groups of lines being consistently low- and high-yielding in both cropping seasons (GY_L_ and GY_H_, respectively). Significance level: *** p < 0.001.

### 3.3. Determinants of grain yield differences in selected lines

GY variation within the GY_L_ and GY_H_ lines was totally unrelated to time to anthesis (Fig. 6a). Regression across all lines, i.e. considering both groups, did show a negative relationship (Fig. 6a) with the lines from sub-group GY_L_ slightly later than that of GY_H_ (Fig. 6b). However, the influence of this difference on GY between the two sub-groups would have been negligible for several reasons. Firstly, the overall GY variation explained by time to anthesis variation was relatively small (c. 35%). Secondly, the groups show extensive overlap to the extent that many GY_H_ lines have the same time to anthesis that many GY_L_ lines (Fig. 6a) still having substantially higher yields (Fig. 5). In fact, only few lines in each sub-group account for the significance of the difference in time to anthesis between the two yielding categories. Finally, and in relation to that distribution, the average time to anthesis between GY_L_ and GY_H_ lines was only slightly different (70°C d, equivalent to c. 3 d; Fig. 6b) and that would hardly explain the large difference in average yield (>300 g m^−2^; Fig. 5).

**Figure 6.**
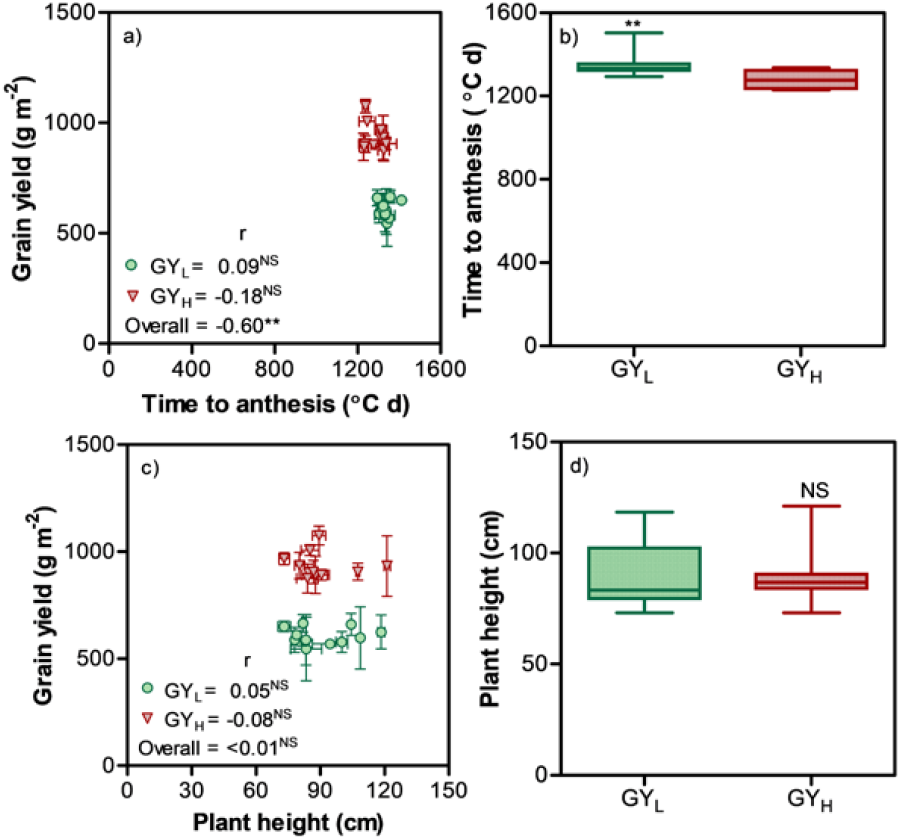
Relationships between yield and either time to anthesis (a) or plant height (c) for the selected sub-groups of low and high yielding lines (GY_L_ and GY_H_, respectively); and boxplots describing the variation in these traits for the two sub-groups (b, d). Coefficients of correlations are shown for each sub-group individually and for the overall data across both sub-groups. Significance level: ** p<0.01; NS=non-significant.

Regarding plant height, although relatively more variable than time to anthesis, the lack of influence of this trait on GY was even more clear, as the relationships were not significant both within and across yielding sub-groups (Fig. 6c), and the range of variation in plant height between the two sub-group contrasting in GY was totally overlapped implying that across them the difference in height was not significant (Fig. 6d).

The fact that neither of these two traits had a relevant role is important as we were interested in identifying likely traits responsible for differences in GY of elite material other than time to anthesis and plant height. And this lack of relevance was also evident if the analysis were made with the whole sub-set of 231 lines (see above and Fig. 4d, e).

Considering the two major GY components, it was clear that variations in GY were almost solely explained by GN (Fig. 7). Considering the variation in GY within sub-groups, it was better explained by those in GN though the coefficient of correlation was non-significant for both GY_L_ and GY_H_ (Fig. 7a) it was still higher than the coefficient of correlation for GY and AGW within any of the two sub-groups (Fig. 7c); In addition, it is also true that the highest yielding line in the GYH sub-group had intermediate GN but the highest AGW within that sub-group (Fig. 7a and c). But most importantly when trying to determine the reasons why the GY_H_ sub-group out-yielded the GY_L_ sub-group, GN was a far more robust determinant of GY than AGW considering all lines across both sub-groups (cf. Fig. 7a and 7c, where it can be seen that more than 70% of the overall variation in GY was related to that in GN, while this percentage was less than 40% when considering AGW).

**Figure 7.**
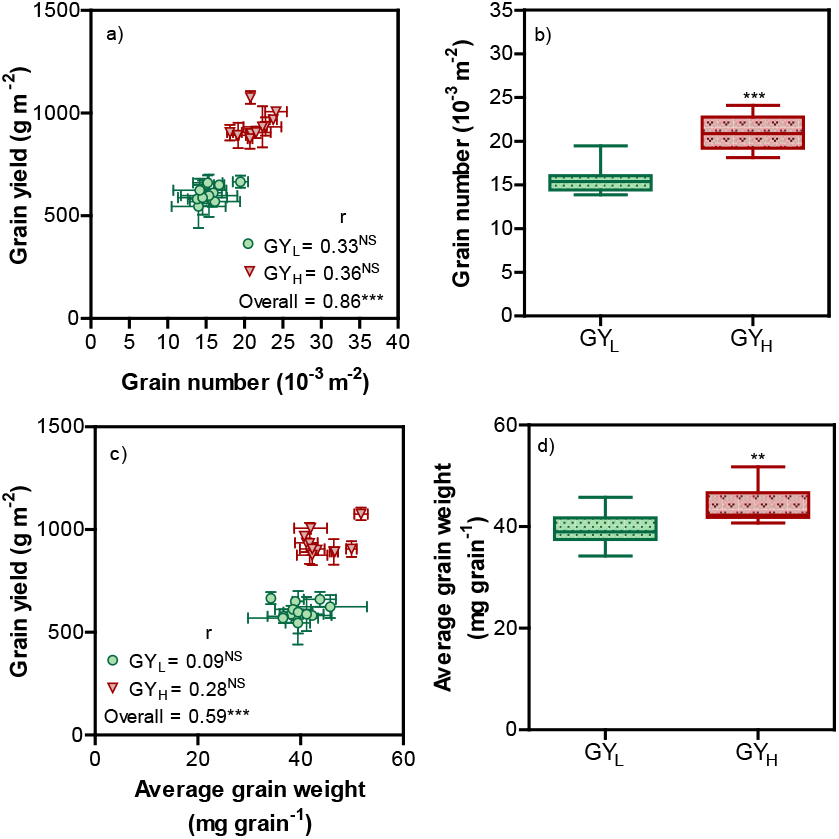
Relationships between grain yield and its two major components: grain number (a) and average grain weight (b) for the selected sub-groups of low and high yielding lines (GY_L_ and GY_H_, respectively); and boxplots describing the variation in these traits for the two sub-groups (b, d). Coefficients of correlations are shown for each sub-group individually and for the overall data across both sub-groups. Significance level: *** p<0.001; NS=non-significant.

Finally, whilst there was virtually no overlap between the ranges of GN of GY_L_ and GY_H_ lines, which differed as groups significantly (in average GY_H_ lines had almost 40% more grains m^−2^ than GY_L_ lines; Fig. 7b), AGW of GY_L_ and GY_H_ lines display noticeable overlapping (in average GY_H_ lines had c. 10% heavier grains than GY_L_ lines; Fig. 7d).

The relationship between these GY components were clearly negative within each of the two sub-groups (Supplementary Fig. S4). However, this did not represent complete compensation as in both sub-groups increasing GN increased GY (Fig. 7a). Furthermore, the negative relationship lost significance when all lines were considered together as the GY_L_ lines did have fewer grains but not so consistently lighter (Supplementary Fig. S4).

The proposed focus on GN was reinforced by our analysis of the variation within the sub-set of 231 lines in each of the two seasons (Supplementary Fig. S5). GN was significantly related to GY in both seasons and GY was also significantly related to AGW although only in CS2 and the magnitude of the association was marginal in absolute terms and negligible compared with that of GN (cf. Supplementary Fig. S5a and b). In both seasons the two major GY components were negatively related across all lines but again this negative relationship did not result in a clear compensation (Supplementary Fig. S5c).

Thus, to understand the traits responsible for the yield advantage of GY_H_ over the GY_L_ lines, it is GN which requires further dissection.

### 3.4. Physiological components of grain number

Physiological components of GN, SDWa and FE, explained part of GN variation (Fig. 8). However, their relative relevance seemed to vary with the type of comparison. When comparing the differences across GY_L_ and GY_H_ lines it seemed that SDWa was most important as the overall relationship was significant (Fig. 8a) and the GY_L_ lines exhibited significantly lower values than those of GY_H_ (Fig. 8b), whereas this trait was unrelated to GN within each of the two sub-groups (Fig. 8a). In general, the contribution of FE to differences in GN was lower than that of SDWa. Although GN in GY_L_ lines was better explained by FE than SDWa, this was not true for the differences in GN within the GY_H_ lines (cf. Fig. 8a and c). Furthermore, the explanatory capacity of FE decreased noticeably when considering the relationship across all the lines (Fig. 8c) and FE was not significantly different between the GY_H_ and GY_L_ lines (Fig. 8d). Indeed, there was a clear negative relationship between the two physiological determinants of GN, mainly driven by the different genotypes within each of the two sub-groups (Supplementary Fig. S6), with the GY_H_ lines being displaced to the right as a result of their overall higher SDWa (i.e. the increase in SDWa of the GY_H_ lines compared to the GY_L_ lines did not bring about any compensation in FE; Figs. 8b, d and S6).

**Figure 8.**
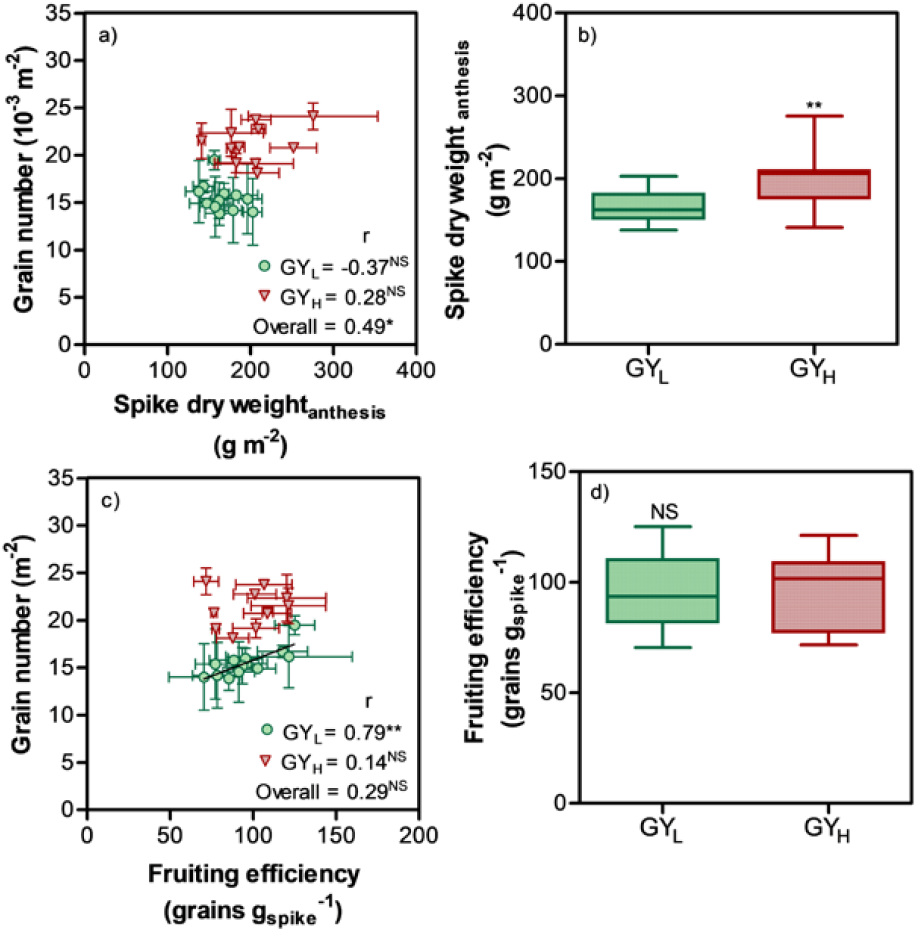
Relationships between grain number and two of its physiological determinants: spike dry weight at anthesis (a) and fruiting efficiency (c) for the selected sub-groups of low and high yielding lines (GY_L_ and GY_H_, respectively). Box plots describing variability for these traits in two sub-groups (b, d). Coefficients of correlations are shown for each sub-group individually and for the overall data across both sub-groups. Significance level: * p<0.05; ** p<0.01; NS=non-significant.

Again, should we have focused the analysis in the comparison in sub-set of 231 lines, the outcome would have been similar. Regardless of the trade-off exhibited by FE and SDWa (Supplementary Fig. S7c), both similarly influenced GN across all lines in each of the two cropping seasons (Supplementary Fig. S7a and b).

The negative relationship between GN and AGW (Supplementary Fig. S5c) is mirrored by the negative relationship between AGW and FE. It is the latter which best explains differences in GN within the GY sub-groups (Supplementary Fig. S8a). This indirect association with genotypes having higher FE tending to have lower AGW was only significant in GY_L_ lines but not in GY_H_ and it was also evident when analysis in sub-set of the 231 lines was considered (Supplementary Fig. S8b).

## 4. Discussion

The present work aimed at identifying traits responsible for GY differences among lines derived from crosses of elite germplasm, beyond the effects of time to anthesis and plant height. Although not considering time to anthesis, we were interested in the partitioning of phenological time in the duration of phases occurring before and after TS. In line with previous research (Halloran and Pennell, 1982; Slafer, 2003; Whitechurch et al., 2007) there was a large variation in the two pre-anthesis phases; and there was clear negative relationship between these two phases. This confirms that it would be possible to lengthen the duration of the LRP at the expense of shortening the duration of the phase from sowing to TS (Miralles et al., 2000; Scarth et al., 1985; Slafer et al., 2001). In this context, importance of these observations rests on the hypothesis that a longer LRP would accommodate increased partitioning of biomass to the growing spike (González et al., 2005b; Miralles et al., 2000; Reynolds et al., 2005; Slafer et al., 2005) with the beneficial knock on effect of increased floret survival and final grain number (Ferrante et al., 2013; Sadras and Slafer, 2012), given all other things are constant. However, the relationships between GY and duration of LRP were positive but rather weak, implying that within this experimental material the duration of this phase was not the most critical trait determining yield differences among lines (as also recently seen in Australia; Zhang et al., 2019).

GN was the main component explaining variations in GY pointing us towards a prioritisation of this trait to explore future genetic gains (Slafer et al., 2014), while not suggesting that maximising AGW is unimportant in the ultimate high yielding varieties produced by breeders (as illustrated by the fact that within the selected lines used for the more detailed characterisation the highest yielding lines had a distinctly higher AGW than the others). To plan for the optimisation of both traits it is important to increase understanding of their negative correlation, which was observed here, as in so many previous studies (Miralles and Slafer, 1995; Siddique et al., 1989a; Slafer et al., 2014). A key question is whether the AGW/GN negative relationship is due to competition for carbohydrates during grain filling. Should there be competition among grains, increasing GN might result in a kind of zero-sum game, with GY not changing significantly. Although, this kind of interpretation of a negative relationship seems quite intuitive, there is good evidence for the less intuitive scenario in which grains do not compete for resources during grain filling. It follows that the source of the trade-off lies elsewhere and is probably not the consequence of a competitive dynamic (Acreche and Slafer, 2006; Miralles and Slafer, 1995). This means that further increases in GN are critical in bringing about major improvements in GY (Fischer, 2011; Reynolds et al., 2012; Sanchez-Garcia et al., 2013; Slafer et al., 2014). These extra grains will have access to sufficient resources for filling as supported by studies from source-sink manipulations during grain filling in which grain growth is unresponsive (Abbate et al., 1997; Borrás et al., 2004b; M. P. Reynolds et al., 2005; Serrago et al., 2013 and references quoted there in; Slafer and Savin, 1994) showing that an excess of assimilates are available at this developmental stage (e.g. Bingham et al., 2007; Borrill et al., 2015); although some exceptions can be found and only for particular seasons under extremely high yielding conditions (e.g. Lynch et al., 2017). In other words, that the crop is rather conservative at the time of establishing GN (Sadras and Slafer, 2012), which would be the reason for the differences in plasticity of GN and AGW (being GN determination strongly responsive to source strength and AGW relatively unresponsive).

The current study did not involve source-sink interventions like defoliations, shading, de-graining, or thinning the plots and so on, nonetheless a quantitative analysis can be used to estimate whether strong source limitation during the grain filling period was at play. For this purpose, we related AGW to the post-anthesis accumulation of crop growth on a per grain basis (i.e. the ratio between total above ground crop dry weight accumulated from anthesis to physiological maturity and GN). This showed that AGW differences between lines were hardly due to severe source limitations in the low AGW genotypes (Fig. 9a). Firstly, there was no clear trend to reduce AGW with reductions in post-anthesis whole crop growth per grain. Secondly, the distributions of the data-points in the figure would be compatible with no source-limitation. Almost all the lines in GY_H_ were very close to the 1:1 line indicating that the grain growth had more than enough resources: cases in which data points are below the 1:1 line would have never being source-limited to the level that even some of the growth produced after anthesis was allocated to other sinks, while cases in which grain weight exceeded the post-anthesis crop growth per grain would still be sink-limited, as post-anthesis growth is only part of the source available to fill the grains (part of the demand of the growing grains can be satisfied by remobilisation of pre-anthesis reserves). This is also true for the data-points from GY_L_ that fell at the left side of the 1:1 line (Fig. 9a). To have a scale that can be more readily compared with the literature we transformed the independent variable into a percentage of AGW (Fig. 9b). Calculated as percent difference between AGW and post-anthesis crop growth per grain with respect to AGW. A negative value means the percentage of GY that was allocated to other organs (i.e. clearly unrealised yield potential due to post-anthesis sink limitation), and positive values represent the percentage of AGW that has been realised thanks to the remobilisation of pre-anthesis reserves. The dotted line at 35% indicates a practical limit up to which developing grains can access translocated pre-anthesis reserves derived from Savin and Slafer (1991). This is a rather conservative figure as there have been examples in the literature where up to 50% of the final grain weight was contributed by translocation of reserves accumulated before anthesis (e.g. Borrás et al., 2004; Gent, 1994) which produced an elegant demonstration of the fact that only with a large contribution of pre-anthesis reserves to grain growth the observed AGW would have been possible. In that work it was estimated that, depending on the cultivar and season, up to 55% of the final AGW could be contributed from pre-anthesis reserves and several examples of such large contribution have been observed (see examples in the review by Blum, 1998).

**Figure 9.**
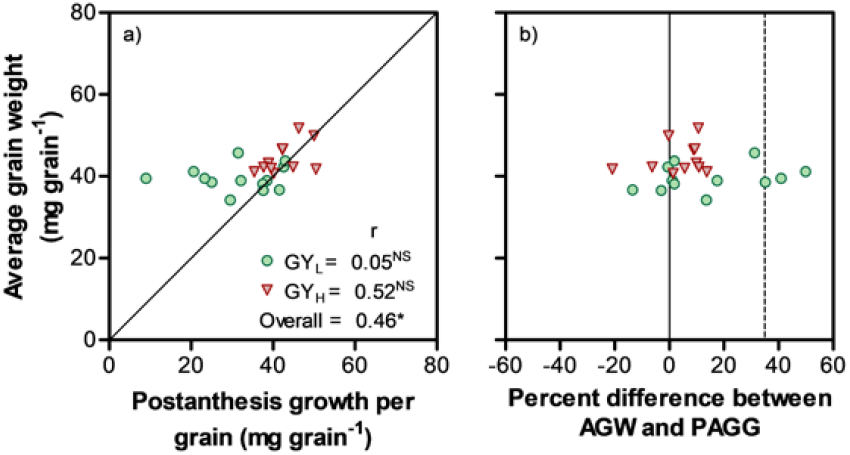
Relationships between average grain weight and either (i) post-anthesis growth per grain (PAGG) in absolute (a), or ii) percent difference between AGW and PAGG with respect to AGW (b) for the selected sub-groups of low and high yielding lines (GY_L_ and GY_H_, respectively). Coefficients of correlations are shown for each sub-group individually and for the overall data across both sub-groups. Significance level: * p<0.05; NS=non-significant. Plain lines represent the situation when AGW was equal to post-anthesis growth per grain. The dotted line represents a 35% contribution from pre-anthesis reserves to final grain weight, which is more than a highly likely contribution that can be expected (Austin *et al*., 1980 in barley; Savin and Slafer, 1991 in wheat).

Furthermore, there was a relationship between GY and total dry weight (at physiological maturity) explaining the genotypic GY differences within and across the GY_L_ and GY_H_ lines (Fig. 10a). The most frequent interpretation of this relationship would be that lines with improved growth capacity produced more grains that, when filled, resulted in a proportionally larger GY. However, this seems not to be the most likely explanation in this case. Looking at the differences in total accumulated dry weight from sowing to anthesis (TDWa; Fig. 10b) and from anthesis to maturity i.e., cumulative growth after anthesis (Fig. 10c), it seems that the more common cause-consequence hypothesis can be inverted to reach an interpretation that is at least as likely. Indeed, there was only a marginal difference in TDWa between GY_L_ and GY_H_ lines, with a large overlap in this trait between lines of the two sub-groups (Fig. 10b), while the difference became relevant in post-anthesis growth (Fig. 10c). Thus, it may well be that the physiological basis for the higher GY of the GY_H_ lines is more efficient translation of pre-anthesis growth into GN. These lines increased the sink strength during grain filling lowering the extent of sink limitation. This, in turn, would reduce the down regulation of post-anthesis canopy photosynthesis (that has been shown to exist due to insufficient sink demand in different conditions; e.g. Serrago et al., 2013) driving the improved crop growth during grain filling. This would be in line with previous studies showing that higher GN would increase post-anthesis growth (Acreche and Slafer, 2009; Reynolds et al., 2005), through its positive effect on canopy photosynthesis.

**Figure 10.**
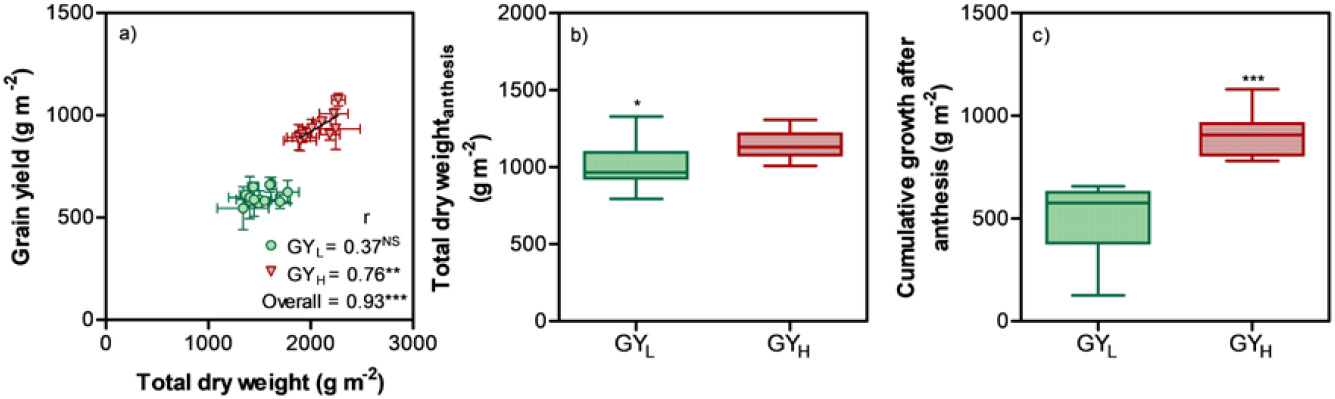
Relation between total dry weight (at maturity) and grain yield (a); box plots showing variations in total dry weight at anthesis (b) and cumulative growth after anthesis (c) for the selected sub-groups of low and high yielding lines (GY_L_ and GY_H_, respectively). Coefficients of correlations are shown for each sub-group individually and for the overall data across both sub-groups. Significance level: ** p<0.01; *** p < 0.001; NS=non-significant.

Both physiological components of GN, SDWa and FE, seemed to have been relevant to improve GY, which is in line with recent results from Australia in a study combining many commercial cultivars, elite lines and a MAGIC population (Zhang et al., 2019). As lines did not differ much in TDWa their differences in SDWa implies a better dry matter partitioning to the juvenile spike growing immediately before anthesis in high GY_H_ lines. This is critical because wheat GY is clearly source-limited just prior to anthesis (Borrás et al., 2004; Slafer and Savin, 1994) and SDWa is critical in determining post-anthesis sink strength (Fischer, 2011; Slafer, 2003). This is because the development of labile florets in the juvenile spikes immediately before anthesis is highly sensitive to allocation of resources (Ferrante et al., 2013, 2010; Fischer, 1985; González et al., 2005a; Siddique et al., 1989b; Slafer et al., 2015), which in turn is the mechanistic basis for the strong and consistent relationship between GN (or number of fertile florets) and SDWa (Fischer, 1985 and a plethora of papers confirming this relationships in different background conditions, in response to various different treatments). In the past, breeding has improved GY through increasing GN exploiting this mechanism. Specifically, modern semi-dwarf cultivars out yielded their older traditional height (tall) predecessors due to an improved dry matter partitioning to the spike before anthesis (e.g. Brooking and Kirby, 1981; Calderini et al., 1995; Fischer and Stockman, 1986; J E Flintham et al., 1997; Miralles et al., 1998; Shearman et al., 2005; Siddique et al., 1989a; Slafer and Andrade, 1993). But these gains were achieved through plant height reduction. In the present study we showed that there is opportunity to further improve partitioning of dry matter to the spike beyond reductions in plant height (Foulkes et al., 2011) that would be instrumental to further improve GY through reducing the degree of sink-limitation during the post-anthesis phases of development. A recent paper by Rivera-Amado et al. (2019) clearly illustrates how this further improvement in pre-anthesis partitioning to juvenile spikes would be possible. The other trait that can help in reducing the sink-limitation during grain filling is FE (e.g. (Slafer et al., 2015). In this study FE explained part of the GN differences within the segregants from elite parents. Although we observed trade-off between SDWa and FE that had been also reported before (e.g. Ferrante et al., 2012; Lázaro and Abbate, 2012), it seemed feasible to identify genotypes with best combinations of both traits maximising GN, and therefore crossing parents with high SDWa and high FE could increase the likelihood of transgressive segregation for GN (and GY).

## Acknowledgement

Funding for the experimental work was partly provided by projects AGL2015-69595-R, from the Spanish Research Agency (AEI), and IWYP25FP, from the International Wheat Yield Partnership (IWYP). We thank Semillas Batlle for their technical support in maintenance of field experiments. We are grateful to the team of laboratory of crop physiology for assisting with field cum laboratory work, and Prof. Ignacio Romagosa for helping with the experimental design and statistical analysis. PB held a pre-doctoral research contract from the agency for Management of University and Research (AGAUR) from the *Generalitat de Catalunya*.

