## Supplementary figures for "Genotypic differences in wheat yield determinants within a NAM population based on elite parents"

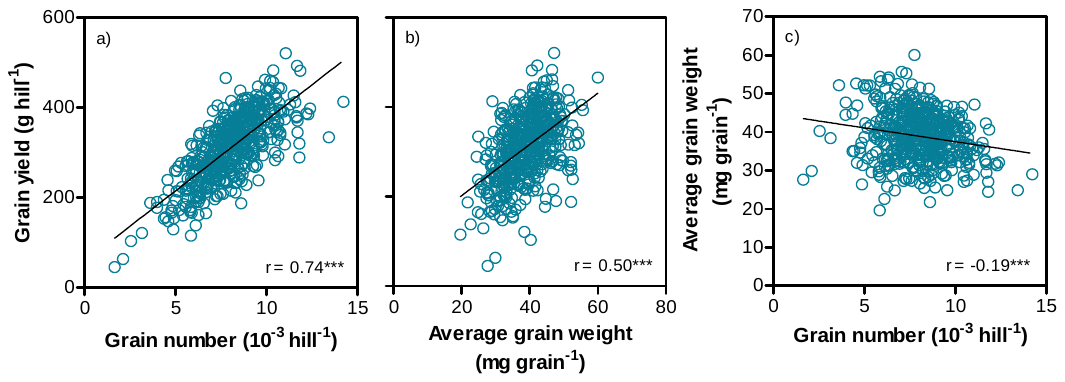

**Supplementary Figure S1. Relationships between hill-plot yield and its two major components (a, b) and between them (c) in the experiment carried out in Cd. Obregón (NW Mexico) in 2015-16 for the 493 lines selected for having a narrow range of time to anthesis, in which yield was determined. Significance level: *** p < 0.001.**

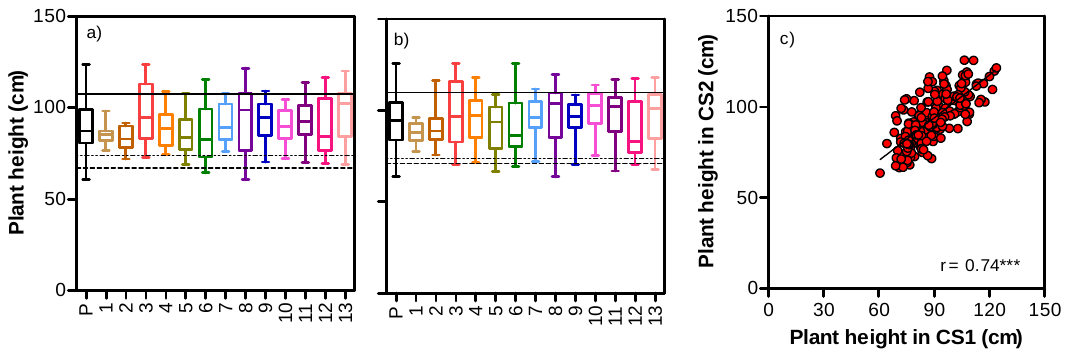

**Supplementary Figure S2. Boxplots showing variability for plant height within the whole population (P_W_) and within families (13 bi-parental crosses) along with three checks Paragon (solid line), Paledor (dotted line) and Garcia (dashed line) in the first (CS1, a) and second cropping season (CS2, b), and consistency for plant height over the two cropping seasons (c). Significance level: *** p < 0.001.**

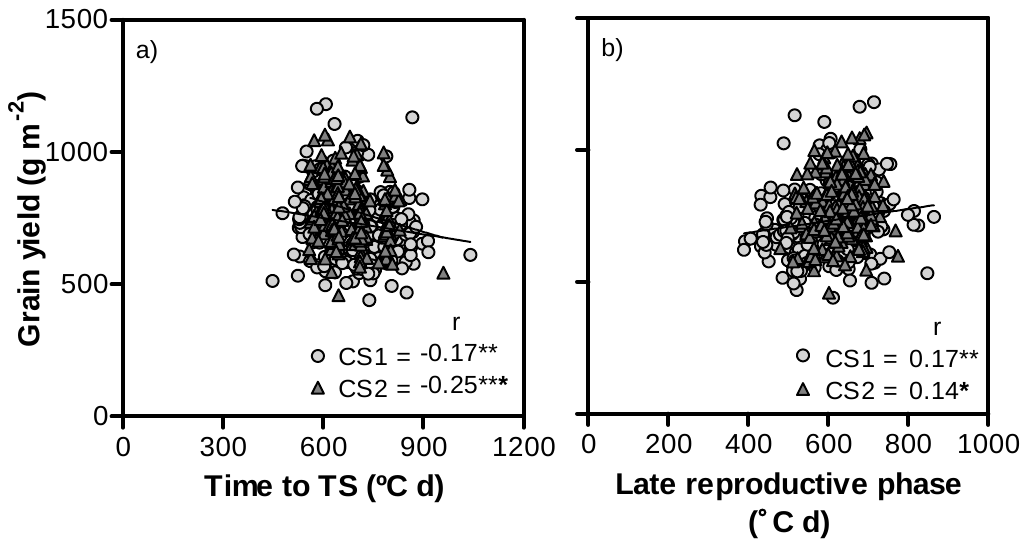

**Supplementary Figure S3. Relationships between grain yield and two component phases of time to anthesis: time from sowing to terminal spikelet (a), and time from then to anthesis, the late reproductive phase (b) in the selected sub-set of 231 lines. Significance level: * p < 0.05; ** p < 0.01; *** p < 0.001.**

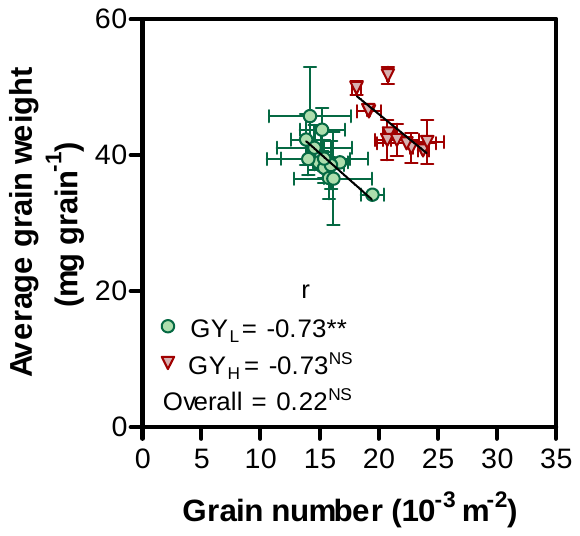

**Supplementary Figure S4. Relationship between the two components of grain yield, average grain weight and grain number, for the two sub-groups of low and high yielding lines (GY_L_ and GY_H_, respectively). Coefficients of correlations are shown for each sub-group individually and for the overall data across both sub-groups. Significance level: ** p < 0.01; NS= non-significant.**

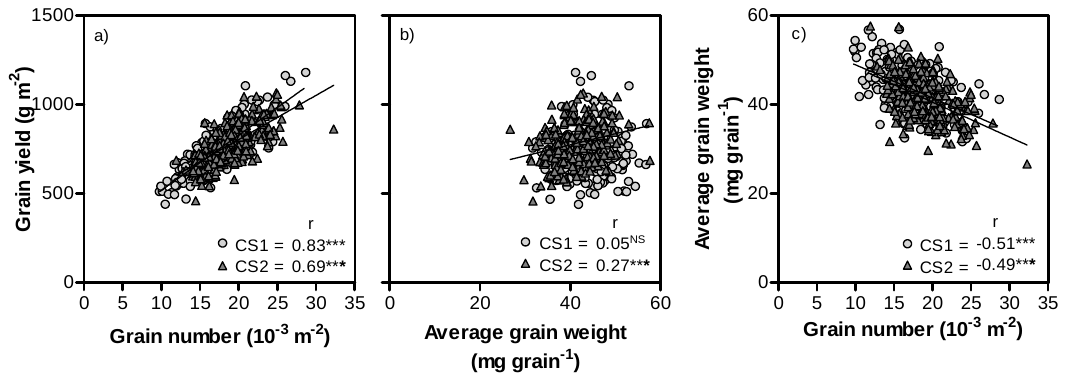

**Supplementary Figure S5. Relationships between grain yield and its components: grain number (a) and average grain weight (b); and between them (c) in the sub-set of 231 lines. Significance level: *** p < 0.001; NS= non-significant.**

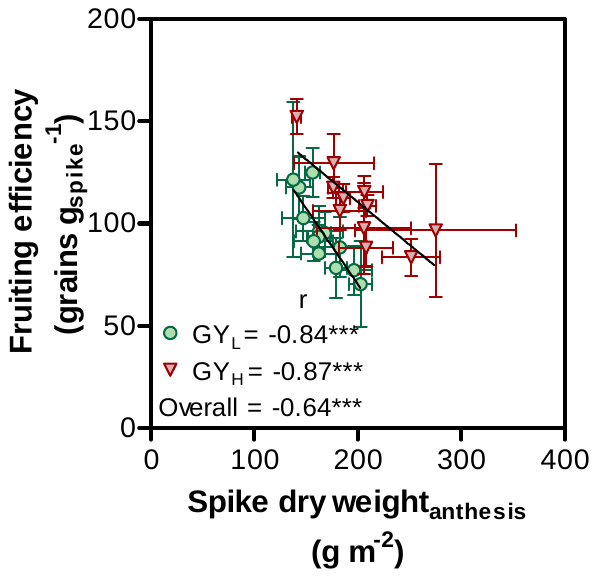

**Supplementary Figure S6. Relationship between spike dry weight at anthesis and fruiting efficiency for the two sub-groups of low and high yielding lines (GY_L_ and GY_H_, respectively). Coefficients of correlations are shown for each sub-group individually and for the overall data across both sub-groups. Significance level: *** p < 0.001.**

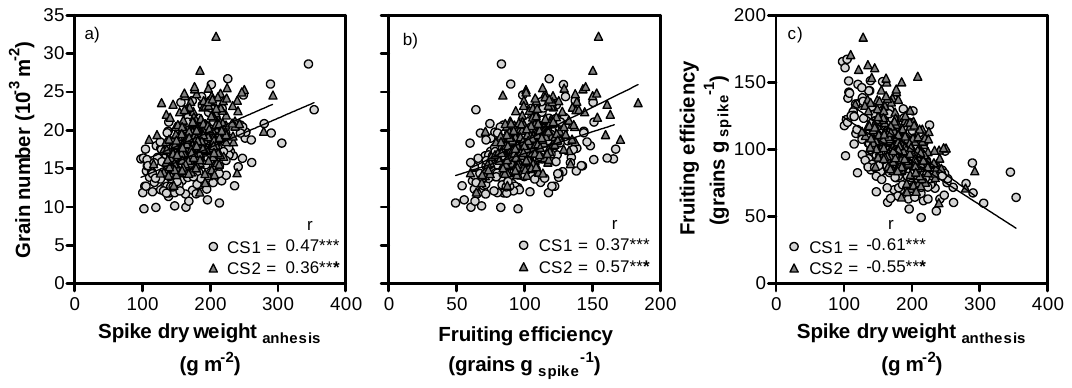

**Supplementary Figure S7. Relations between grain number and two of its physiological determinants: spike dry weight at anthesis (a) and fruiting efficiency (b); relation between spike dry weight and fruiting efficiency (c) in the sub-set of 231 lines. Significance level: *** p < 0.001.**

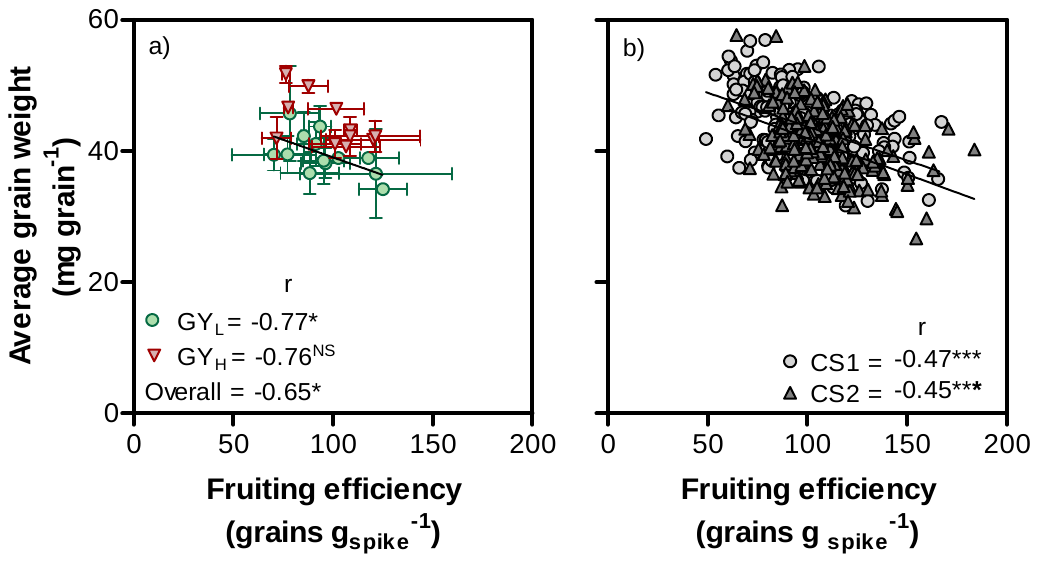

**Supplementary Figure S8. Relation between fruiting efficiency and average grain weight in selected sub-groups of low and high yielding lines (GY_L_ and GY_H_, respectively) with coefficients of correlations are shown for each sub-group individually and for the overall data across both sub-groups (a) and in sub-set of 231 lines (b). Significance level: * p < 0.05; *** p < 0.001; NS= non-significant.**
